# The structures of protein kinase A in complex with CFTR: mechanisms of phosphorylation and reversible activation

**DOI:** 10.1101/2024.05.28.596263

**Authors:** Karol Fiedorczuk, Iordan Iordanov, Csaba Mihályi, András Szöllősi, László Csanády, Jue Chen

## Abstract

Protein kinase A (PKA) is a key regulator of cellular functions by selectively phosphorylating numerous substrates, including ion channels, enzymes, and transcription factors. It has long served as a model system for understanding the eukaryotic kinases. Using cryo-electron microscopy, we present complex structures of the PKA catalytic subunit (PKA-C) bound to a full-length protein substrate, the cystic fibrosis transmembrane conductance regulator (CFTR) – an ion channel vital to human health. CFTR gating requires phosphorylation of its regulatory (R) domain. Unphosphorylated CFTR engages PKA-C at two locations, establishing two “catalytic stations” near to, but not directly involving, the R domain. This configuration, coupled with the conformational flexibility of the R domain, permits transient interactions of the eleven spatially separated phosphorylation sites. Furthermore, we determined two structures of the open-pore CFTR stabilized by PKA-C, providing a molecular basis for understanding ‘reversible activation’, whereby PKA-C stimulates CFTR function through simple binding.

## Introduction

Protein kinases comprise approximately 2% of the human genome (1) and regulate the activities of more than 30% of cellular proteins (2). Consequently, kinases have become common drug targets for a wide range of diseases, with more than 70 drugs targeting kinases currently approved for clinical use (3, 4).

The cyclic AMP (cAMP)-dependent protein kinase, also known as protein kinase A (PKA), was the first kinase to be purified (5) and has become a prototype of the eukaryotic protein kinase family (reviewed (6); (7)). The inactive PKA holoenzyme is a heterotetramer of two regulatory (R) and two catalytic (C) subunits. The binding of cAMP relieves the inhibition imposed by the R subunits, permitting the C subunits to phosphorylate a large number of target proteins. The PKA-C consists of a small N-terminal lobe (N-lobe) and a larger C-terminal lobe (C-lobe) joined by a flexible linker. The active site cleft, located between the two lobes, contains several highly conserved motifs that catalyze phosphoryl transfer from ATP to a serine or threonine of the protein substrate. Since the publication of the first PKA-C structure in 1991 (8), over three hundred structures of PKA-C bound to various nucleotides, peptides, and drugs have been reported. In addition, the structure of the holoenzyme (PKAC2:RIIβ2) was determined by X-ray crystallography, highlighting the dual role of RIIβ as both a substrate and an inhibitor of the catalytic subunit (9). However, thus far, no structure of PKA-C in complex with a downstream target protein is available. The absence of information on how PKA interacts with protein substrates is a significant gap in our knowledge. As highlighted by Taylor and Kornev, “protein kinases do not work on small peptides in cells; they phosphorylate proteins” (10).

An example of a protein activated by PKA is the cystic fibrosis transmembrane conductance regulator (CFTR), an epithelial anion channel crucial for salt-water balance of the lung, intestine, pancreas, and sweat duct (11). CFTR loss-of-function mutations cause cystic fibrosis (CF) (12), and CFTR hyperactivation due to abnormally high PKA activities underlies diarrhea in cholera (13)) and cyst growth in autosomal dominant polycystic kidney disease (ADPKD) (14). PKA-C regulates CFTR through two distinct mechanisms. CFTR gating (opening and closure of the pore) is under tight control by its regulatory (R) domain, a mostly unstructured segment containing 11 consensus PKA-C phosphorylation sites (15). The unphosphorylated R domain inhibits gating by preventing the conformational changes necessary for pore opening (16). Phosphorylation by PKA releases the R domain from its inhibitory position (17), permitting ATP-binding induced dimerization of its nucleotide binding domains (NBDs) to open, and ATP hydrolysis to close the channel. A secondary activation by PKA involves simple binding of PKA-C to CFTR, which enhances the channel current independent of the catalytic activity (18). The process, termed reversible activation, is distinct from the irreversible activation achieved through phosphorylation.

Small-molecule drugs that bind and stabilize the CFTR channel in the open-pore conformation (19, 20) are currently used to treat CF patients (21). An alternative approach to enhance or dampen CFTR activity – depending on disease condition – would be to manipulate the extent of CFTR phosphorylation using molecules that modulate the affinity of PKA to CFTR. However, developing such drugs requires detailed structural information about how PKA interacts with CFTR.

In this work, we used cryo-electron microscopy (cryo-EM) to uncover the molecular interactions between PKA-C and full-length CFTR, both with and without phosphorylation. These structures advance our understanding of how PKA-C recognizes and activates an ion channel with multiple phosphorylation sites.

## Results and Discussion

### PKA-C phosphorylates CFTR by creating multiple catalytic stations

Previous mass spectrometry and NMR studies have collectively detected ten phosphoserines in the R domain and one in NBD1 (22–25). Modelling the positions of these residues onto the structure of CFTR shows that they are broadly scattered (Figure 1A), raising the question of how PKA-C acts on the multiple sites across such a large spatial distribution.

**Figure 1.**
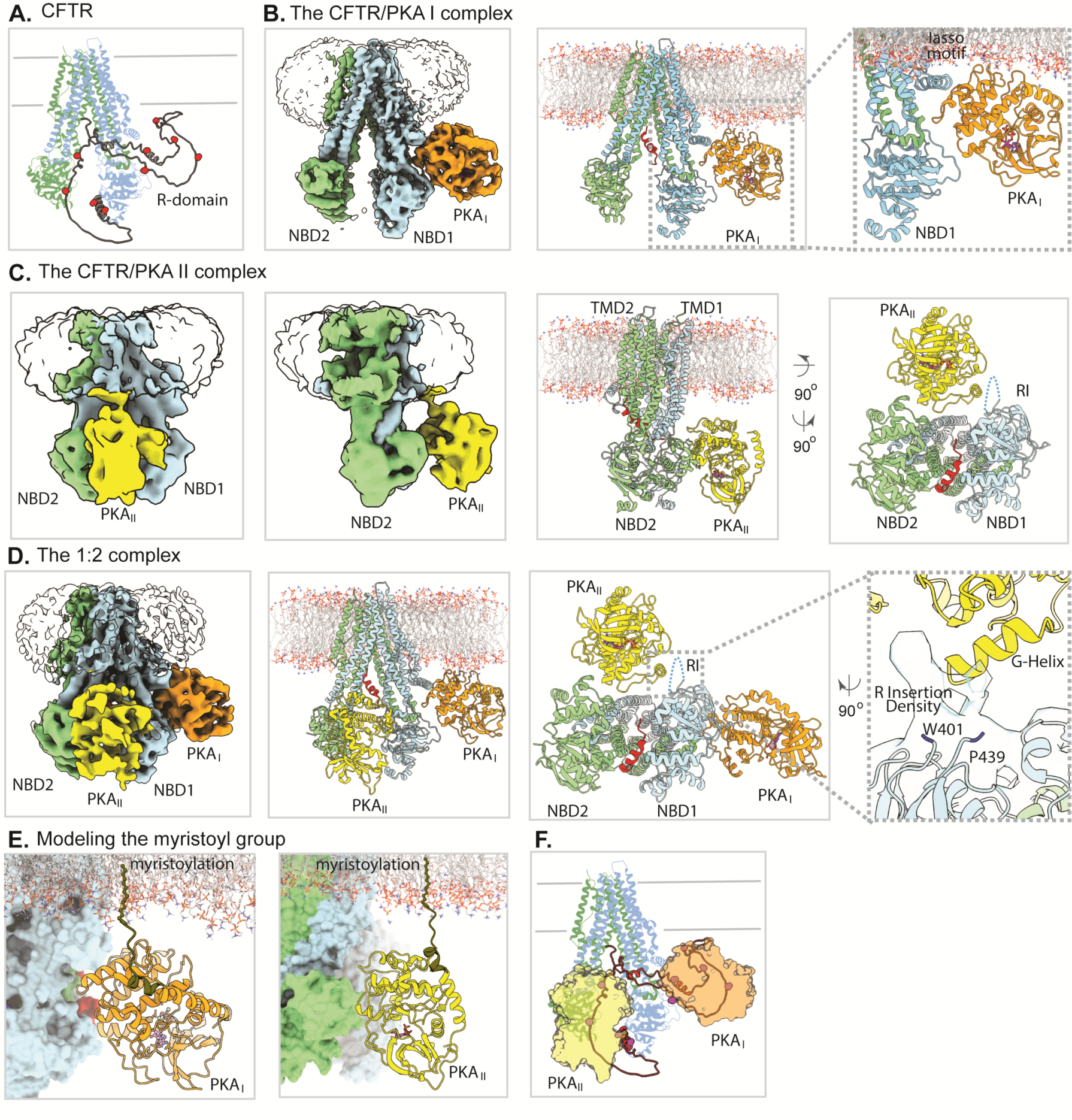
PKA-C phosphorylates CFTR by creating multiple catalytic stations. **(A)** Structure (PDBID: 5UAK) of dephosphorylated CFTR (TMD1-NBD1, *blue*; TMD2-NBD2, *green*), with the R domain modelled by Alphafold (*gray ribbon*). *Red spheres* represent phosphorylation sites at positions 660, 670, 686, 700, 712, 737, 753, 768, 795, and 813. **(B)** EM density (*left*) and model (*center*) of the 1:1 complex of CFTR with PKA_I_ (*orange)*. PKA_I_ docks against CFTR’s lasso helix 2 and the NBD1-TMD1 linker (*right*). **(C)** Two views of the EM density (*left two panels*) and model (*right two panels*) of the 1:1 complex of CFTR with PKA_II_ (*yellow)*. PKA_II_ docks against the helical subdomain of NBD2 and the region around the RI of NBD1 (*right*). **(D)** EM density (*left*) and model (*center, right*) of the 1:2 complex of CFTR with PKA-C. PKA_I_ (*orange)* and PKA_II_ (*yellow)* bind at two distinct sites on CFTR (*right*). **(E)** Modeling the PKA-C N terminal residues 1-14 (*brown*) places the myristoyl groups of both PKA_I_ (*orange)* and PKA_II_ (*yellow)* into appropriate positions for membrane insertion. **(F)** All phosphorylation sites can be accessed by PKA-C parked to either site I or site II.

To address this question, we used cryo-EM to investigate the interactions between PKA-C and dephosphorylated CFTR. The sample, comprising 30 µM recombinant human CFTR purified in digitonin, 250 µM PKA-C purified from bovine heart (bPKA-C), and 2 mM AMP-PNP—known to bind and stabilize PKA-C but not support phosphorylation—is very heterogenous. Three distinct CFTR/PKA-C complexes were observed including two 1:1 complexes with a PKA-C molecule bound at different locations (termed site I and site II, respectively) and a 1:2 complex with both sites occupied (Figure 1B-D, and Figure S1A). Despite rigorous data processing, the final resolutions were somewhat limited: 3.8 Å for the PKA_I_-bound complex, 9.7 Å for the PKA_II_-bound complex, and 6.0 Å for the 1:2 complex.

The conformation of CFTR in all three complexes is similar, and closely resembles that of the dephosphorylated CFTR in the absence of PKA (16). Briefly, the two transmembrane domains (TMDs) form a domain-swapped inverted “V” configuration, the two nucleotide-binding domains (NBDs) are separated, and density for the R domain is largely missing. The absence of well-defined density for the R domain indicates that the unstructured nature of the R domain persists even in the presence of PKA-C.

The two PKA-C binding sites are spatially separated, appearing nearly orthogonal to each other when viewed from the cytosolic side of the membrane (Figure 1D). PKA_I_ binds on one side of CFTR, between the N-terminal lasso motif and NBD1 (Figure 1B). PKA_II_ docks between the two NBDs, interacting mainly with the helical subdomain of NBD2, and is also within van der Waals contact with an unassigned density protruding from NBD1 (Figure 1C). This density is connected to residues W401 and P439 in NBD1, and as such likely represents the R insertion (RI) in which the phosphorylation site S422 resides. In the NBD-separated conformation, the two NBDs are inherently flexible, leading to the lower resolution structures of the PKA_II_-containing complexes.

Despite their distinct positions on CFTR, both PKA-C molecules are oriented with their N-terminal ends projecting towards the membrane (Figure 1E). No density was observed for the N-terminal 14 residues of either PKA_I_ or PKA_II_. Modeling this region (Figure 1E) suggests that the N-terminal myristoyl groups for both PKA-C molecules are within the range to be inserted into the membrane. These observations align with data showing the marked enhancement of PKA-C efficiency by the myristoyl group (see accompanying paper), suggesting that the myristoyl group facilitates the membrane partitioning of PKA-C, thereby elevating its local concentration. While this has only been demonstrated for CFTR, it is likely that, for other membrane protein substrates, the N-myristoyl group also plays a crucial role in enhancing the activities of PKA-C at the membrane surface.

The presence of two distinct PKA-C binding sites on CFTR provides a plausible explanation for how the eleven widely dispersed PKA sites can all be phosphorylated. Mapping these positions onto the 1:2 complex structure indicates that nearly all phosphorylation sites are in proximity to one of the two PKA-C molecules (Figure 1F). Because the R domain is largely unstructured and highly flexible, individual peptide segments containing a phosphorylation site (i.e, peptide substrates) can reach into one of the catalytic sites.

To examine the relative abundance of the three complexes, we sorted the entire dataset of 3.3 million particles into different structural classes through 3D heterogeneous refinement (Figure S1B). The results revealed a comparable distribution of the three CFTR/PKA-C complexes, indicating that PKA-C binds to the two sites independently and with similar affinities. These findings provide a plausible explanation for earlier observations that multiple sites in intact CFTR seem to undergo phosphorylation simultaneously (26, 27).

Lastly, both PKA _I_ and PKA _II_ interact with CFTR through a region away from the active site cleft. No density corresponding to peptides was observed in the catalytic site, despite clear density for AMP-PNP in PKA_I_. These data indicate that PKA-C acts not by binding tightly to the R domain, but rather by docking onto other regions of CFTR. The “parked” PKA-C molecules serve as catalytic stations, allowing peptide substrates to enter the active site, undergo phosphorylation, and then exit to create space for the phosphorylation of a different peptide substrate.

### PKA-C stabilizes phosphorylated CFTR in an open-pore conformation

In addition to phosphorylating the R domain, PKA-C also activates CFTR through simple binding. Robust current stimulation was observed upon addition of PKA-C to already phosphorylated CFTR (18). To elucidate the structural basis of this activity, we determined the structure of PKA-C in complex with fully-phosphorylated CFTR carrying the ATP-hydrolysis-disrupting E1371Q mutation, in the presence of 3 mM ATP (Figure 2 and S2). In contrast to the dephosphorylated CFTR, we observed only a single 1:1 complex wherein a PKA-C binds at site I of an NBD-dimerized CFTR. The absence of PKA_II_ is a result of the conformational changes triggered by phosphorylation and ATP binding. As the two NBDs approach to form the closed dimer, the structure becomes incompatible with PKA_II_ binding. In contrast, the structure of site I remains identical in both conformations of CFTR, capable of engaging PKA_I_ regardless of the phosphorylation state.

**Figure 2.**
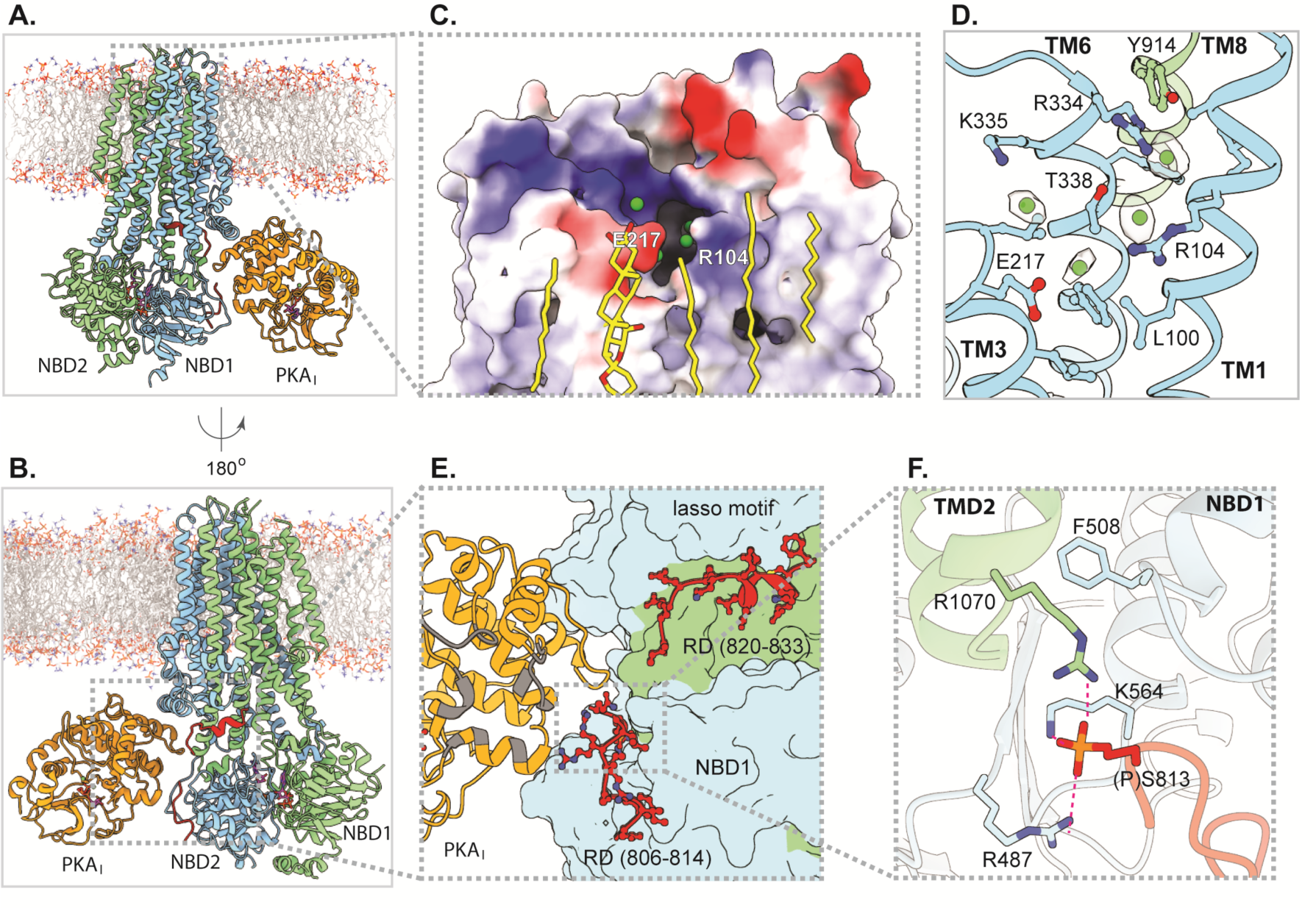
PKA-C stabilizes phosphorylated CFTR in a channel-open conformation. **(A-B)** Two views of the phosphorylated E1371Q CFTR in complex with PKA-C. *Red ribbon* represents resolved R-domain segments (residues 806-814, 820-833) of CFTR, PKA-C (*orange*) binds to site I. *Gray dotted boxes* highlight regions magnified in panels (C) and (E). **(C)** Positive surface electrostatics of the outer pore exit, with putative chloride ions (*green spheres*) and tightly bound lipids (*yellow sticks*). **(D)** Putative chloride ion densities in and around the selectivity filter and side chains involved in anion coordination. **(E)** Model of resolved R-domain segment (*red sticks*) bound to the external surface of CFTR’s TM11-TM12 and NBD1. Interaction surface of PKA-C with PKI(6–22)-amide is highlighted in *gray*. *Gray dotted box* marks region magnified in panel (F). **(F)** Phosphoserine 813 is buried in a cavity at the NBD1/TMD interface, anchored to basic side chains.

The structure, determined at 2.8 Å resolution, unveils two previously unobserved features of CFTR. First, the luminal exit of the pore (Figure 2A, *gray dotted box*) is better defined through a string of ion-like densities connecting the selectivity filter and the extracellular space (Figure 2C, D). The selectivity filter in CFTR is positioned at the extracellular ends of TM 1, 6, and 8, where a dehydrated chloride ion is coordinated by highly conserved residues G103, R334, F337, T338, and Y914 (28). In the PKA-stabilized structure, the density for the chloride ion within the selectivity filter is prominent, along with several other densities that may represent chloride ions exiting the pore (Figure 2C,D). These densities are within range to interact with R104, R334, and K335— residues previously shown to influence ion conduction (29–33). The observation of a continuous ion pathway provides support for the earlier proposition that the NBD-dimerized E1371Q structure represents the conductive state of CFTR (28).

Secondly, two regions within the R domain revealed well-defined density, permitting us to assign the amino acid registry for residues 806-814 and 820-833 (Figure 2E and S2B). The visualization of phosphoserine S813 has important mechanistic implications, as many studies underscore its critical role in CFTR activation (27, 34–36). The phosphate group of S813 inserts into a cavity at the NBD1/TMD interface, engaging in electrostatic interactions with R487 and K564 of NBD1, as well as R1070 of TMD2, substitutions of which are associated with cystic fibrosis (Figure 2F). As discussed earlier (37), molecular contacts at the NBD1/TMD interface are comparatively weaker than those at the NBD2/TMD interface, making this region susceptible to destabilization by mutations. Thirteen cystic fibrosis-causing mutations, including the F508 deletion, target the NBD1/TMD interface (37). The structural data indicate that phosphorylation of S813 likely reinforces the NBD1/TMD interface, thereby playing a pivotal role in CFTR function. In the structures of a related protein, the yeast glutathione transporter Ycf1, a segment of its R domain is positioned similarly to the phosphorylated CFTR R domain (38, 39). However, unlike in CFTR, where the phosphoserine 813 is buried at the NBD1/TMD interface, in Ycf1 the phosphate group on the equivalent residue is exposed to the solvent (39).

### PKA-C binding opens CFTR pore even without phosphorylation

CFTR activation via PKA-C binding occurs not only with phosphorylated CFTR but is also evident in situations where R-domain phosphorylation is not feasible (18). To understand how PKA-C opens the dephosphorylated CFTR channel, we determined the cryo-EM structure of dephosphorylated CFTR (30 μM) in the presence of bovine PKA-C (250 μM) and the ATP analog N^6^-(2-phenylethyl)-ATP (P-ATP, 1 mM), which supports CFTR channel gating with an apparent affinity (K_d_) of 2 μM (40) but does not bind to PKA-C (18, 41). We also included AMP-PNP (1 mM), which has a higher affinity for PKA-C (K_d_ = 35 μM) than for CFTR (K_d_ = 300 μM) (42, 43). Under such conditions a 1:1 complex of CFTR/PKA-C was determined to 3.5 Å resolution (Figure 3 and S3). Consistent with the apparent affinities and concentrations of the nucleotides used, P-ATP molecules are bound to CFTR whereas the nucleotide binding pocket in PKA-C is occupied by AMP-PNP (Figure 3A and Figure 3D-F).

**Figure 3.**
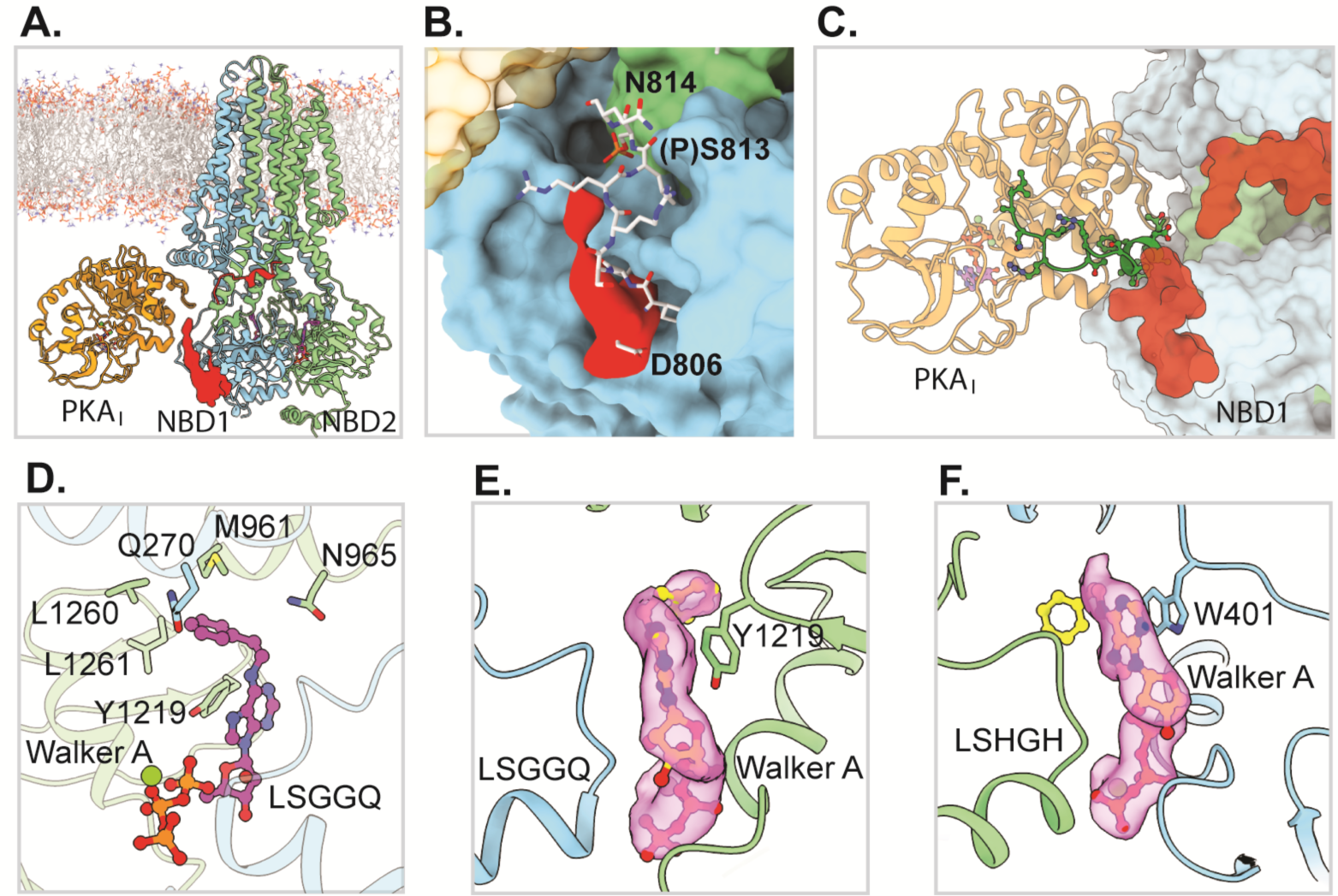
PKA-C binding opens dephosphorylated CFTR. **(A)** Structure of the WT, dephosphorylated CFTR in complex with PKA-C obtained in the presence of P-ATP and AMP-PNP. **(B)** The dephosphorylated R domain segment 806-813 in P-ATP-bound structure (*red surface*) is positioned similarly to the phosphorylated R domain (*white sticks*) observed in complex with ATP and PKA-C. **(C)** Structure of PKI(6–22)-amide (*green*; from PDBID: 2GFC) aligned with phosphorylated ATP-bound CFTR:PKA-C complex. CFTR is shown as a surface, PKA-C and PKI as ribbon diagram. Note steric clash between PKI N-terminal helix and the R-domain (*red surface*). **(D)** Molecular details of P-ATP (shown in ball-and-stick representation) in the canonical ATP-binding site. Side chains of residues in NBD2, CH2, and CH3 interacting with the phenylethyl group are shown in sticks. (**E-F)** EM densities (*magenta surfaces*) of P-ATP in the canonical (E) and degenerate (F) nucleotide-binding sites; flanking Walker A and signature motifs are shown as ribbons.

Although PKA-C can use neither AMP-PNP nor P-ATP to phosphorylate the R domain, the structure closely resembles that of the phosphorylated CFTR/PKA-C complex (RMSD of 0.80 Å over 1517 Cα positions). The density for the unphosphorylated R segment 806-813 was less resolved, but its position closely resembles that of the phosphorylated version (Figure 3B). These data indicate that PKA-C activates the unphosphorylated CFTR by stabilizing the same open-pore structure as in the phosphorylated channel, with the R segments wrapping around TM11-TM12 and NBD1. When the R domain is phosphorylated, the phosphoserine 813 enhances this configuration by anchoring the R-segment to NBD1 (Figure 2F). Interference with that effect may explain inhibition of reversible CFTR activation by PKI(6–22)-amide (accompanying manuscript), as structural alignment reveals steric clashes between the N-terminal helix of PKI and the R-domain segment on the outer face of NBD1 (Figure 3C).

Taken together, the structures of PKA-C complexed with either phosphorylated or dephosphorylated CFTR illustrate that reversible activation is achieved through an allosteric mechanism, not by direct binding of the R domain to segregate it away from the inhibitory position. It is likely that PKA-C, by interacting mainly with NBD1 and the lasso motif, lowers the free energy of the NBD-dimerized state, thereby promoting channel opening in the presence of ATP (or P-ATP) for both phosphorylated and dephosphorylated CFTR. Furthermore, the effect of PKI(6–22)-amide suggests that this stabilizing effect may also require positioning of the R-domain loop in the conformation observed in our structures.

The structure of the P-ATP-bound CFTR also provides a framework for understanding the activity of P-ATP, which gates the CFTR channel with kinetic features distinct from ATP. The higher affinity of P-ATP for stimulating channel opening rate (40) reflects tighter binding of P-ATP to the consensus site (44). The longer openings (bursts) under both hydrolytic and non-hydrolytic conditions (40) have been ascribed to P-ATP bound at the degenerate site (44–46). The two P-ATP molecules bound at the NBD interface adopt different conformations. Density for the P-ATP in the consensus site is well defined, exhibiting a kinked conformation with the phenyl and adenine rings forming an angle of approximately 100° (Figure 3D-E). The phenylethyl group reclines towards NBD2 and forms multiple interactions with residues within NBD2 (Y1219, L1260) and coupling helices (CH) 2 and 3 (Q270, M961) (Figure 3D). These molecular interactions, absent for ATP, explain the higher affinity of P-ATP for the consensus site. The NBD2-CH2-CH3 unit moves like a rigid body during the CFTR gating cycle (47), preserving the phenylethyl binding pocket in both the channel open and closed (interburst) states. In contrast, in the degenerate site density for the phenylethyl group of P-ATP is less well resolved (Figure 3F), suggesting that it is flexibly exposed at the NBD interface.

### Perturbations of the site I interface impair CFTR activation by PKA-C

PKA-C binds to both dephosphorylated and phosphorylated CFTR at site I, suggesting its critical role in both reversible and irreversible activation mechanisms. The molecular details of site I are best defined in the 2.8 Å structure obtained with phosphorylated CFTR (E1371Q) (Figure 2 and 4A). PKA_I_ interacts with CFTR through a narrow interface, burying ∼720 Å^2^ per subunit. The relative small buried surface area suggests low affinity protein-protein interaction, underscoring the necessity of having the N-terminal myristoyl group to enrich and orient PKA in the membrane. Two regions are involved at the PKA_I_/CFTR interface (Figure 4). One region (Figure 4B) is formed through electrostatic interactions between the H-helix and the preceding G-H loop of PKA-C (segment 254-274) and the CFTR lasso helix 2 (segment 46-63). Four acidic residues implicated in channel activation (48), namely D47, E51, E54, and D58, are aligned along one face of lasso helix 2, forming a predominantly negative surface. This surface interacts via diffuse electrostatic forces with a positive surface patch on PKA-C lined by the basic residues K254, R256, K266, and R270. The strongest ionic interaction forms between the side chains of D58 of CFTR and R270 of PKA-C, which approach each other to approximately 2.8 Å.

**Figure 4.**
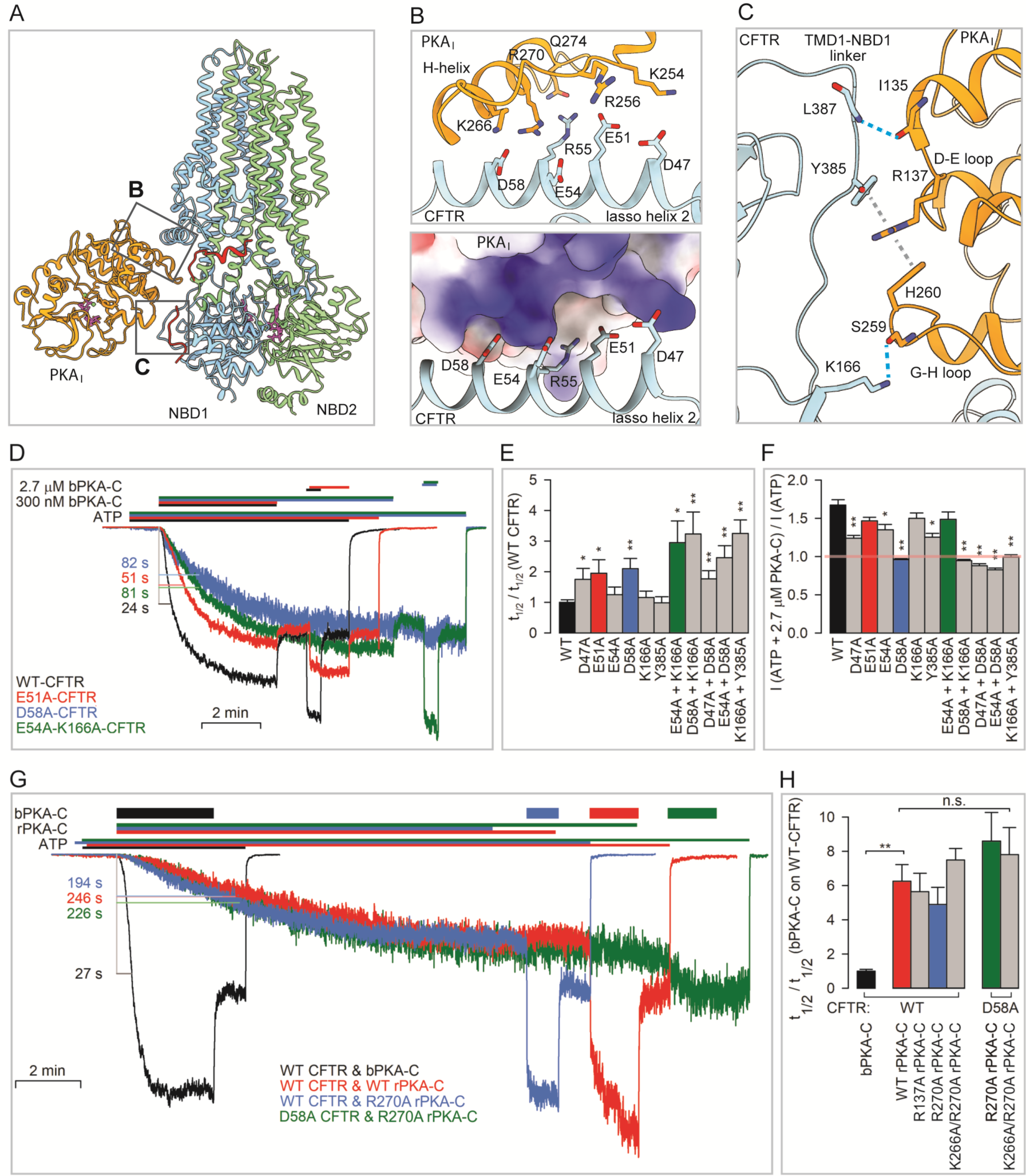
Molecular details and functional relevance of PKA site I. **(A)** Ribbon diagram of the 1:1 CFTR/PKA-C complex determined in the presence of MgATP. Boxes highlight regions magnified in panels (B) and (C). **(B)** Zoomed-in view of interface 1 with charged side chains shown as sticks (top) and electrostatics of the PKA-C surface patch in contact with acidic side chains of the CFTR lasso helix 2 (bottom). **(C)** Zoomed-in view of interface 2 with interacting residues shown as sticks. **(D)** Inside-out macropatch currents of WT and mutant (color coded) CFTR channels activated by 300 nM, and then briefly exposed to 2.7 μM, bovine PKA-C (bPKA-C) in 2 mM ATP. Color coding of the bars that identify exposure times to compounds follows that of the current traces. Currents are normalized to their steady-state amplitudes in 2 mM ATP, following full phosphorylation by 2.7 μM bPKA-C. L-bars and numbers illustrate the time required for half-maximal current activation (t_1/2_). **(E)** Activation half-times (t_1/2_) in 300 nM bPKA-C for various mutant CFTR constructs, normalized to t_1/2_ of WT CFTR obtained in parallel experiments in the respective oocyte batches. **(F)** Reversible stimulation by bPKA-C for WT and mutant CFTR constructs, expressed as the ratio of mean steady current in ATP + 2.7 μM PKA-C to that following bPKA-C removal. Colored bars in (E) and (F) identify the four representative constructs shown in (D). **G)** Currents of WT and D58A CFTR channels activated either by 300 nM bPKA-C, or by various rPKA-C constructs, in 2 mM ATP. Trace colors represent particular combinations of a CFTR and a PKA-C construct; color coding of the bars that identify exposure times to compounds follows that of the current traces. Currents are normalized to their steady-state amplitudes in 2 mM ATP, following full phosphorylation by ∼2 minute exposure to 300 nM bPKA-C. L-bars and numbers illustrate the time required for half-maximal current activation (t_1/2_). **(H)** Half-times (t_1/2_) of current activation by various rPKA-C constructs (300 nM) for WT and D58A CFTR, normalized to t_1/2_ of WT CFTR in 300 nM bPKA-C obtained in parallel experiments in the respective oocyte batches. Colored bars in (H) identify the four representative CFTR/PKA-C construct pairs shown in (D). Data in (E), (F), (H) represent mean±SEM, n=10-17, 6-14, 4-8, respectively. Asterisks highlight significant changes relative to the control conditions (*black bars*); p<0.05 (*), p<0.01 (**).

A second region in site I (Figure 4C) is formed by the TMD1-NBD1 linker and the cytosolic ends of TM2 and TM11 of CFTR, which contact the D-E loop (segment 135-137) and G-H loop (segment 256-260) of PKA-C (Figure 4C). The side-chain guanidino group of R137 in PKA-C is sandwiched between the aromatic side chains of H260 in PKA-C and Y385 in the CFTR TMD1-NBD1 linker (Figure 4C, *gray dashed lines*). In addition to the strong arginine-tyrosine cation-pi interaction, multiple H-bonds (Figure 4C, *cyan dashed lines*) provide further stability. Despite decades of extensive studies, the structural elements of this interface have not yet been identified to play a functional role in either CFTR or PKA-C.

To test the functional relevance of site I, we used alanine substitutions in CFTR to perturb the acidic lasso-helix residues D47, E51, E54, and D58 in region 1, and K166 and Y385 in region 2, and quantified mutational effects in inside-out patch-clamp recordings (Figure 4D). Effects on channel phosphorylation (irreversible activation) were estimated by comparing the rates of macroscopic current activation upon exposure to 300 nM bPKA-C and quantified through activation half-times (t_1/2_, Figure 4D, L-bars). Consistent with the large area of the site I interface, individual CFTR mutations only modestly affected the rate of current activation, but a significant slowing was evident for single mutants D58A, E51A, and D47A which prolonged t_1/2_ by approximately twofold compared to WT (Figure 4E). Furthermore, combinations of single mutations typically showed additive effects. Thus, although single mutations E54A, K166A, and Y385A caused no significant slowing, for the E54A/K166A or K166A/Y385A double mutants t_1/2_ was again significantly prolonged (Figure 4E).

Mutational effects on reversible channel activation, caused by PKA-C binding, were quantified through the fractional current increase upon a brief exposure of pre-phosphorylated channels to 2.7 μM bPKA-C (Figures 4D, 4F). Reversible activation was slightly but significantly reduced for single mutants D47A, E54A, and Y385A, but entirely abolished by the D58A mutation (Figure 4D, *blue trace*; Figure 4F, *blue bar*). Reversible activation was similarly eliminated by a combination of the two “weak” mutations K166A and Y385A in interface 2 (Figure 4F, *right*). These results, consistent with structural observations, suggest that reversible CFTR stimulation is mediated by the PKA-C bound in site I.

To address the effects of perturbing the site I interfaces from the PKA-C side, the wild-type, R270A, K266A/R270A (interface 1), and R137A (interface 2) bovine PKA-C sequences were expressed as recombinant proteins in *E. coli* (rPKA-C). All constructs were affinity-purified to homogeneity (Figure S4A-B) and displayed catalytic activities towards the soluble substrate peptide kemptide comparable to each other and to bPKA-C (k_cat_ ∼7-9 s^-1^, Figure S4C-H). The efficiencies of WT and mutant rPKA-C proteins towards CFTR channel activation were studied in inside-out patch recordings (Figure 4G). As shown in the accompanying manuscript, due to lack of the N-myristoyl group in the recombinant protein the rate of CFTR current activation by rPKA-C is severalfold slower compared to bPKA-C (Figure 4G, *red* vs. *black trace*). Indeed, t_1/2_ was found ∼6-fold prolonged (Figure 4H, *red* vs. *black bar*). Interestingly, that slow rate of current activation was not further affected by any of the three PKA-C mutations (Figure 4G, *blue trace*; Figure 4H, *2nd group* of *bars*). Moreover, when activated by any rPKA-C protein, no significant difference between t_1/2_ for WT or D58A CFTR currents was apparent (Figure 4G, *green* vs. *blue trace*; Figure 4H, *3rd* vs. *2nd group* of *bars*). These results suggest that the nonmyristylated rPKA-C may bind to CFTR differently as compared to myristylated bPKA-C. That conclusion also explains the complete lack of reversible CFTR stimulation by rPKA-C (Figure 4G, *colored traces*; also see accompanying manuscript).

### Implications for PKA signaling

The structures of the CFTR/PKA-C complex offer for the first time a molecular description of how PKA interacts with a protein substrate. The core structures of PKA-C (residues 13-337 and 343-351) within the various CFTR complexes are very similar, and closely resemble that of isolated PKA-C stabilized by Mg^2+^ and nucleotide (Figure 5A-B). Among the three autophosphorylation sites that regulate the activity of PKA-C (S10, T197, and S338), S10 is in a region not resolved in the structure. The other two sites both have clear densities for the phosphate moiety, indicating that the bPKA-C purified from the native source is phosphorylated and catalytically active.

**Figure 5.**
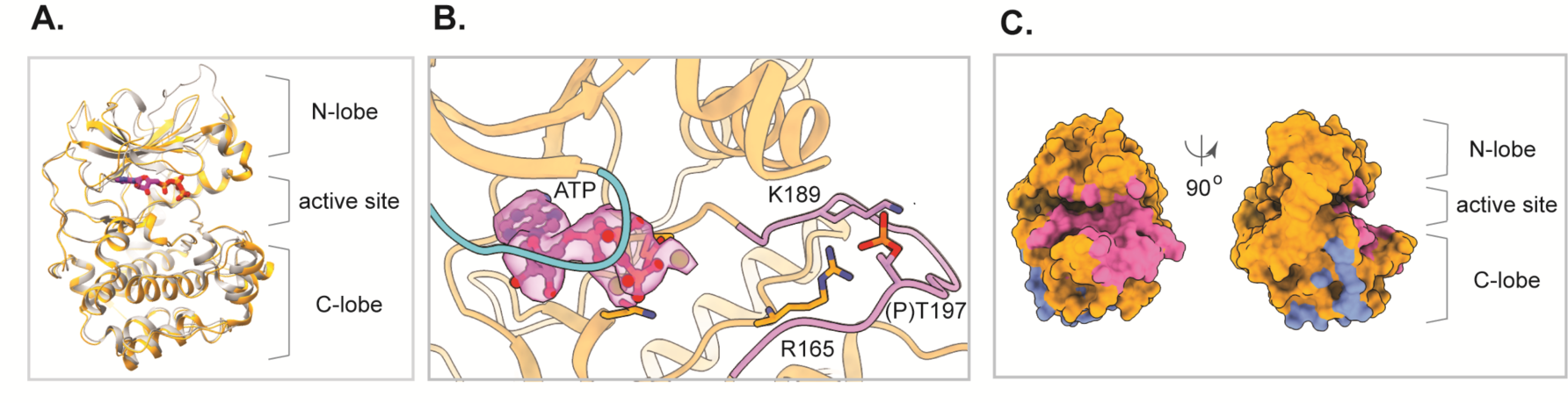
Structural features of PKA-C in site I. **(A)** Structure of ATP-bound bPKA-C from the activated 1:1 complex (*orange*) aligned with the crystal structure of mouse PKA-C (*gray*) bound with AMPPNP and the substrate peptide SP20 (PDBID: 4DG0). ATP bound to bPKA-C is shown as sticks. AMPPNP and SP20 were omitted for clarity. **(B)** Zoomed-in view of the bPKA-C substrate binding cleft from (A), with clear density (*purple surface*) for bound Mg-ATP. The glycine rich loop (*cyan*) and the activation loop (*purple*) are colored for reference. Phosphothreonine 197 and interacting basic side chains are shown as sticks. **(C)** Surface representations of bPKA-C in site I; the orientation on the right roughly matches that in panel (A). Colored surface patches identify interaction surfaces with CFTR (*slate*) and with the PKA-RIIβ regulatory subunit (*magenta*), respectively.

Although phosphoryl transfer takes place between ATP and a short segment of the protein containing the consensus phosphorylation site, peptide substrates on their own bind poorly to PKA-C (49). Consistently, none of the cryo-EM reconstructions reveal any peptide density in the substrate-binding cleft of PKA-C, suggesting a lower affinity for individual peptides compared to the binding sites identified within the context of the full-length CFTR.

Our structural and mutational analyses (Figure 4) highlight the functional significance of the interface between PKA-C and CFTR at site I. At site II, due to the limited cryo-EM resolution, specific side chains engaged at the interface cannot be unambiguously identified. Nonetheless, it is evident from the overall position of PKA_II_ that it also interacts with CFTR through a similar surface as observed for PKA_I_ (Figure 5C).

The interface between PKA-C and CFTR, located within the large C-lobe of PKA-C, is distinct from the surface to which the PKA-R subunit binds (Figure 5C), suggesting that small molecules targeting this interface may modulate CFTR phosphorylation without interfering with the cAMP-dependent regulation of PKA-C. Whether these compounds will impact phosphorylation of other substrate proteins remains uncertain, but to our knowledge no data exist to support a role of this surface in interacting with other proteins.

The structures of CFTR/PKA-C complexes help to resolve a longstanding controversy in the field. While much of the literature indicates that PKA-C dissociates from PKA-R as cAMP levels increase, some studies also suggest that phosphorylation is achieved through a complex form of PKA-C and R subunits (50–53). When aligning one C subunit of the intact (RIIβ)_2_(Cα)_2_ holoenzyme (PDBID: 3TNP) with PKA_I_ on CFTR, the holoenzyme clashes with CFTR, indicating that the holoenzyme must dissociate to interact with CFTR. An alternative hypothesis is that a smaller complex, comprising PKA-C and the cyclic nucleotide binding (CNB) domains of the R subunit, interacts with the substrate protein (52, 53). Mapping the PKA-C/CNB complex onto CFTR would immerse the CNB domains deep into the membrane, an unlikely configuration given the hydrophilic nature of the CNB domains (Figure S5). Based on these analyses, we conclude that phosphorylation of CFTR requires dissociation of the catalytic subunit from the PKA holoenzyme.

Finally, kinase-substrate interactions beyond the active site have been previously described for several other eukaryotic kinases (reviewed in (2, 54)). Our findings reveal that PKA also utilizes this strategy to enhance substrate specificity, as the interactions with individual peptide substrates are inevitably transient. Furthermore, the complexity of CFTR demonstrates the possibility and necessity of having multiple kinase-docking sites to fully phosphorylate a large protein with numerous spatially separated phosphorylation sites. Binding of PKA-C at one of the docking sites also allosterically stabilizes CFTR in a pore-open conformation, providing the additional reversible activation observed in electrophysiological studies.

## ACKNOWLEDGMENTS

We thank M. Ebrahim, H. Ng, and J. Sotiris at Rockefeller’s Evelyn Gruss Lipper Cryo-Electron Microscopy Resource Center for assistance in electron microscopy data collection, Dr. P. Zhang at the National Cancer Institute for discussion and comments on the manuscript. This work is supported by the Howard Hughes Medical Institute (to J.C.) and Cystic Fibrosis Foundation Research Grant CSANAD21G0, National Research, Development and Innovation Fund grant KKP 144199, and EU Horizon 2020 Research and Innovation Program grant 739593 (to L.C).

## AUTHOR CONTRIBUTIONS

K.F. performed all the cryo-EM experiments, analyzed the structures, and prepared structural figures and the corresponding method description. I.I. and A.S. purified bPKA-C and rPKA-C, and performed experiments presented in Figure S4. C.M. and L.C. performed the electrophysiology experiments presented in Figure 4. L.C. and J.C. supervised the study and wrote the manuscript. The authors declare no competing financial interests.

## Methods

### Cell culture

Sf9 cells for baculovirus generation were cultured in Sf-900 II SFM medium (GIBCO) supplemented with 5% FBS and 1% Antibiotic-Antimycotic. HEK293S GnTi^-^ cells used for protein expression were cultured in Freestyle 293 (GIBCO) supplemented with 2% FBS and 1% Antibiotic-Antimycotic.

### CFTR protein Expression and Purification

The methodology for expressing and purifying all CFTR constructs follows protocols outlined in prior studies (37, 64) with some modifications. Plasmids carrying CFTR constructs were used to transfect E. Coli DH10Bac cells (Invitrogen) to produce bacmids. Purified bacmids were used to produce baculoviruses in Sf9 cells. The protein expression was performed in HEK293S GnT-cells, which were infected with baculovirus to 10% final concentration at a density of 2.7×106 cells/mL. 12 hours after transfection, cells were treated with 10 mM sodium butyrate and maintained at 30°C for an additional 48 hours before harvesting. For every cryo-EM experiment, protein purification followed a similar protocol with specific adjustments explained below. Cell solubilization was conducted in a buffer containing 1.2% 2,2-didecylpropane-1,3-bis-b-D-maltopyranoside (LMNG) and 0.24% cholesteryl hemisuccinate (CHS). Subsequently, the cell lysate was clarified by centrifugation. The soluble fraction was applied to GFP nanobody coupled Sepharose Beads (GE Healthcare) and eluted by GFP tag removal using PreScission Protease. After this stage, the wild-type samples underwent de-phosphorylation employing lambda phosphatase, whereas the E1371Q mutant samples were de-phosphorylated with lambda phosphatase or phosphorylated with protein kinase A. The final purification step involved size exclusion chromatography in a buffer containing 0.03% digitonin, 200 mM NaCl, 20 mM HEPES (pH 7.4), 1 mM DTT. The buffer was additionally supplemented with 1 mM ATP for the phosphorylated or 1mM P-ATP for the de-phosphorylated E1371Q mutant.

### Purification of native bovine PKA-C from beef heart (bPKA-C)

The catalytic subunit of protein kinase A was purified from beef heart following a published protocol (65), as described (see accompanying manuscript). Approximately 2.7 kg cleaned myocardium was ground and homogenized in 6 liters of Buffer A (10 mM potassium-phosphate (pH=6.9), 1 mM EDTA) supplemented with 15 mM β-mercaptoethanol (β-ME) and 0.1 mM phenylmethylsulfonyl fluoride (”Buffer A+”), and centrifuged at 14,300 x g for 35 min. The supernatant was mixed with 2 liters of DEAE-sepharose (Merck) pre-equilibrated with Buffer A+ at 6.9, and stirred overnight. The resin was dried, washed 4x with 2 liter Buffer A+ and 1x with 2 liter Buffer A+ supplemented with 50 mM NaCl, and eluted with 4 liters of Buffer A+ supplemented with 500 mM NaCl (Buffer B+). The eluate (∼4.5 liters) was salted out with 390 g/l (NH_4_)_2_SO_4_, and centrifuged at 14,300 x g for 35 min. The pellet was resuspended in 450 ml Buffer A+ and dialysed against 8 liters of Buffer A+ for ∼15 h. The dialysed sample (∼600 ml) was centrifuged at 16,000 x g for 30 min, and the pH of the supernatant adjusted to 6.1 with 1M acetic acid. The sample was mixed with 200 ml CM Sephadex C-50 resin preequilibrated with Buffer A+ (pH=6.1), stirred for 5 minutes and filtered through a B<u>ü</u>chner funnel. The flowthrough was recovered and this treatment repeated 4x at pH=6.1 and then 5x at pH=6.9. The sample was centrifuged at 17,700 x g for 30 min, the supernatant was supplemented with 200 μM cAMP and loaded onto a 20-ml CM Sephadex C-50 column preequilibrated with Buffer A+ (pH=6.9). The column was washed with 2×30 ml Buffer A supplemented with 15 mM β-ME, and eluted with a linear salt gradient (50 ml Buffer A vs. 50 ml of 300 mM potassium phosphate (pH=6.9), 1 mM EDTA, 15 mM β-ME). The PKA-C subunit eluted at ∼150 mM potassium phosphate. Fractions of high purity (>90% by SDS-PAGE; Fig. S1C, *left*) were pooled, and the final concentration determined from the optical density at 280 nm (A_280_; NanoPhotometer P300, Implen GmbH). The entire protocol was carried out at 4-6°C.

### EM sample preparation, data acquisition and map reconstruction

Following size exclusion chromatography, the CFTR (wild-type or E1371Q) sample was concentrated to 5 mg/mL. Catalytic subunit of PKA was buffer exchanged to 200 mM NaCl, 20 mM HEPES (pH 7.4) and 1 mM DTT in the presence of ATP/Mg^2+^ or 4mM AMPPNP/Mg^2+^ depending on the final complex composition and concentrated to 10 mg/mL. Final cryo-EM samples were formed by combining CFTR and PKA-C protein samples in 1:1 (v/v) ratio and concentrating back to the original protein concentration of the components (final concentration of the protein in the samples is ∼5 mg/mL of CFTR and ∼10 mg/mL of PKA). Final nucleotide compositions in the samples were: 2 mM AMPPNP/Mg^2+^ in the de-phosphorylated CFTR/PKA-C sample, 3 mM ATP/Mg^2+^ in the phosphorylated E1371Q CFTR/PKA-C sample or 1 mM P-ATP/Mg^2+^ with 1 mM AMPPNP/Mg^2+^ in the de-phosphorylated E1371Q CFTR/PKA-C. Subsequently, approximately 3 mM fluorinated Fos-choline-8 was introduced to the samples just before freezing onto Quantifoil R0.6/1 300 mesh Cu grids using the Vitrobot Mark IV (FEI). Cryo-EM images were captured utilizing a 300 kV Titian Krios (FEI) equipped with a K3 Summit detector (Gatan) and controlled by SerialEM in super-resolution mode. These images underwent gain reference correction and were binned by 2 before drift correction via MotionCorr (56) to pixel size of 0.676 Å. Contrast transfer function (CTF) estimation was accomplished using GCTF (57). Particle picking was performed automatically by Gautomatch (https://www.mrc-lmb.cam.ac.uk/kzhang/). Further steps, including map reconstruction and resolution estimations, were carried out using RELION 4.0 (59) and CryoSPARC (66). The processing strategy for each dataset was slightly modified to yield optimal results (Figure S1). Generally, the initial 2D classification and 3D classifications as well as CTF refinement and Bayesian polishing were performed in RELION. The final non-uniform refinement was performed in CryoSPARC. For the de-phosporylated CFTR/PKA-C complex particles underwent 2D classification and few rounds of 3D classification that generated initial maps of the CFTR/PKA_I_ and CFTR/PKA-C 1:2 complexes. Next, a model of CFTR/PKA_II_ was generated in UCSF Chimera by partial signal subtraction from the CFTR/PKA-C 1:2 map. Those three maps were used as initial models for the several rounds of CryoSPARC heterogeneous refinement of particles selected in 2D classification in Relion (Figure S1). The final subsets of particles were NU refined in CryoSPARC. For the E1371Q datasets particles underwent several rounds of 2D and 3D classifications followed by the CTF refinement and Bayesian polishing to generate a final stack for particles that were exported to CryoSPARC and refined producing final maps.

### Analysis of particles distribution in the de-phosphorylated CFTR/PKA sample

All the particles extracted from the micrographs of the de-phosphorylated CFTR/PKA dataset were exported to CryoSPARC. These particles underwent heterogeneous refinement using default settings. Initial models for refinement included six maps: CFTR/PKA_I_, CFTR/PKA 1:2, CFTR/PKA_II_, zebrafish CFTR (EMD-8516), noisy CFTR, and a detergent micelle map (Figure S1B). With the exception of the zebrafish CFTR (EMD-8516), all maps represent final or intermediate steps in the processing of the de-phosphorylated CFTR/PKA dataset.

### Model building and refinement

Initial protein models were constructed by fitting published CFTR structures (PDB:5UAK and 6O1V) and AlphaFold 2 PKA models into the cryo-EM maps using UCSF Chimera (67). In the de-phosphorylated CFTR/PKA_II_ and CFTR/PKA-C 1:2 structures, the protein side chains were trimmed due to the limited resolution (Figure S1). These models were then adjusted based on the cryo-EM densities using Coot (68) and refined using PHENIX (61). MolProbity (62) was employed for geometry validation.

### Presentation of structures

Structural figures were generated using UCSF ChimeraX (69).

### Molecular biology

The cDNA of bovine protein kinase A catalytic subunit alpha in pJ411, and of WT CFTR in pGEMHE (see accompanying manuscript), served as templates for introducing point mutations using the QuikChange II XL kit (Agilent Technologies). All constructs were confirmed by automated sequencing (LGC Genomics). The cDNA for CFTR in pGEMHE was linearized using Nhe I (New England Biolabs), transcribed *in vitro* (mMESSAGE mMACHINE T7 Transcription Kit, ThermoFisher Scientific), and purified cRNA stored at - 80 °C.

### Expression and purification of recombinant bovine PKA-C (rPKA-C)

WT and mutant PKA-C in pJ411 was transformed into *E. coli* BL21(DE3) and purified as described (see accompanying manuscript). Colonies were grown at 37°C in 1 liter Luria-Bertani (LB) medium supplemented with 50 μg/ml kanamycin until OD_600_ ∼0.5, and protein expression was induced with 0.1 mM isopropyl-β-D-thiogalactoside (IPTG) overnight at 25°C. Cells were lyzed by sonication in Buffer A (100 mM Tris-HCl (pH=7.5), 150 mM NaCl, 0.1 mM EDTA, 10 mM MgCl_2_, 2 mM DTT) supplemented with protease inhibitors. The cleared supernatant was supplemented with 1.5 mg avidin and loaded onto a 5-ml STREP-Tactin Superflow column (IBA Lifesciences). PKA-C was eluted with Buffer A + 10 mM desthiobiotin (IBA Lifesciences). Protein-rich fractions were concentrated (Vivaspin 6, 10,000 MWCO), passed through a Superdex 200 gel filtration column (GE Healthcare). The main peak fractions (Fig. S4A, *colored chromatograms*) were collected, pooled, quality-checked by SDS-PAGE (Fig. Sxx B), and the final concentration calculated from A_280_.

### Kemptide phosphorylation assay

Kemptide phosphorylation was done as described (18). PKA-C protein (5 nM), TAMRA-kemptide (20 μM; Addexbio Technologies), and Mg-ATP (200 μM) were co-incubated at room temperature for 0-12 minutes in reaction buffer (50 mM HEPES (pH=7.5), 10 mM Mg-acetate, 0.2 mg/mL bovine serum albumin, 5 mM DTT). A 2-μl aliquot of each sample was spotted on a TLC sheet (SIL/UV254, Macherey-Nagel), and the sheet developed in a mixture of n-butanol, pyridine, acetic acid, and water (15:10:3:12 v/v). The relative amounts of dephospho-vs. phospho-kemptide in each sample were quantitated by densitometry (ImageJ), and k_cat_ was estimated from the time required for phosphorylation of 10 μM kemptide (obtained by linear interpolation of bracketing time points) (Fig. S4C-H).

### Xenopus laevis oocyte isolation and injection

*Xenopus laevis* oocytes were isolated following Institutional Animal Care Committee guidelines, injected with 10 ng cRNA, and stored at 18°C (70). Inside-out patch-clamp recordings were obtained 2-3 days after injection.

### Inside-out patch-clamp recordings

Inside-out patch-clamp recordings were done as described (18). Patch pipette solution contained 136 mM NMDG-Cl, 2 mM MgCl_2_, 5 mM HEPES, pH=7.4 with NMDG. Bath solution contained 134 mM NMDG-Cl, 2 mM MgCl_2_, 5 mM HEPES, 0.5 mM EGTA, pH=7.1 with NMDG. MgATP (Merck, A9187) was added from a 400 mM aqueous stock (pH=7.1 with NMDG). Purified rPKA-C and bPKA-C were added from 40-60 μM stock solutions. For the addition of 2.7 μM bPKA-C (Fig. 4D) the ∼150 mM potassium phosphate buffer of the bPKA-C stock was exchanged to 10 mM potassium phosphate using a Hi-TRAP desalting column (GE Healthcare). Thus; in all experiments final phosphate concentration following bPKA-C addition remained <∼1 mM. Recordings were done at 25°C at a membrane potential was -40 mV. The composition of the continuously flowing bath solution could be exchanged (ρ=∼20 ms) using electronic valves (ALA-VM8, ALA Scientific Instruments). Currents were amplified, filtered at 2 kHz (Axopatch 200B, Molecular Devices), digitized at a sampling rate of 10 kHz (Digidata 1322A, Molecular Devices), and recorded to disk (pCLAMP 9, Molecular Devices).

### Electrophysiological data analysis and statistics

CFTR currents were Gaussian-filtered at 50 Hz and baseline-subtracted (pCLAMP 9, Molecular Devices). Mean steady-state currents of fully phosphorylated channels in 2 mM ATP + 2.7 μM bPKA-C were normalized to the mean current observed in the same patch in 2 mM ATP, following bPKA-C removal. Current activation half-times (t_1/2_) report the time required for reaching 50% of the final, steady-state current amplitude. Data represent mean±S.E.M., the number of experiments is indicated in each figure legend. Significances were evaluated using Student’s t-test (*p<0.05; **p<0.01).

---

**Figure S1.**
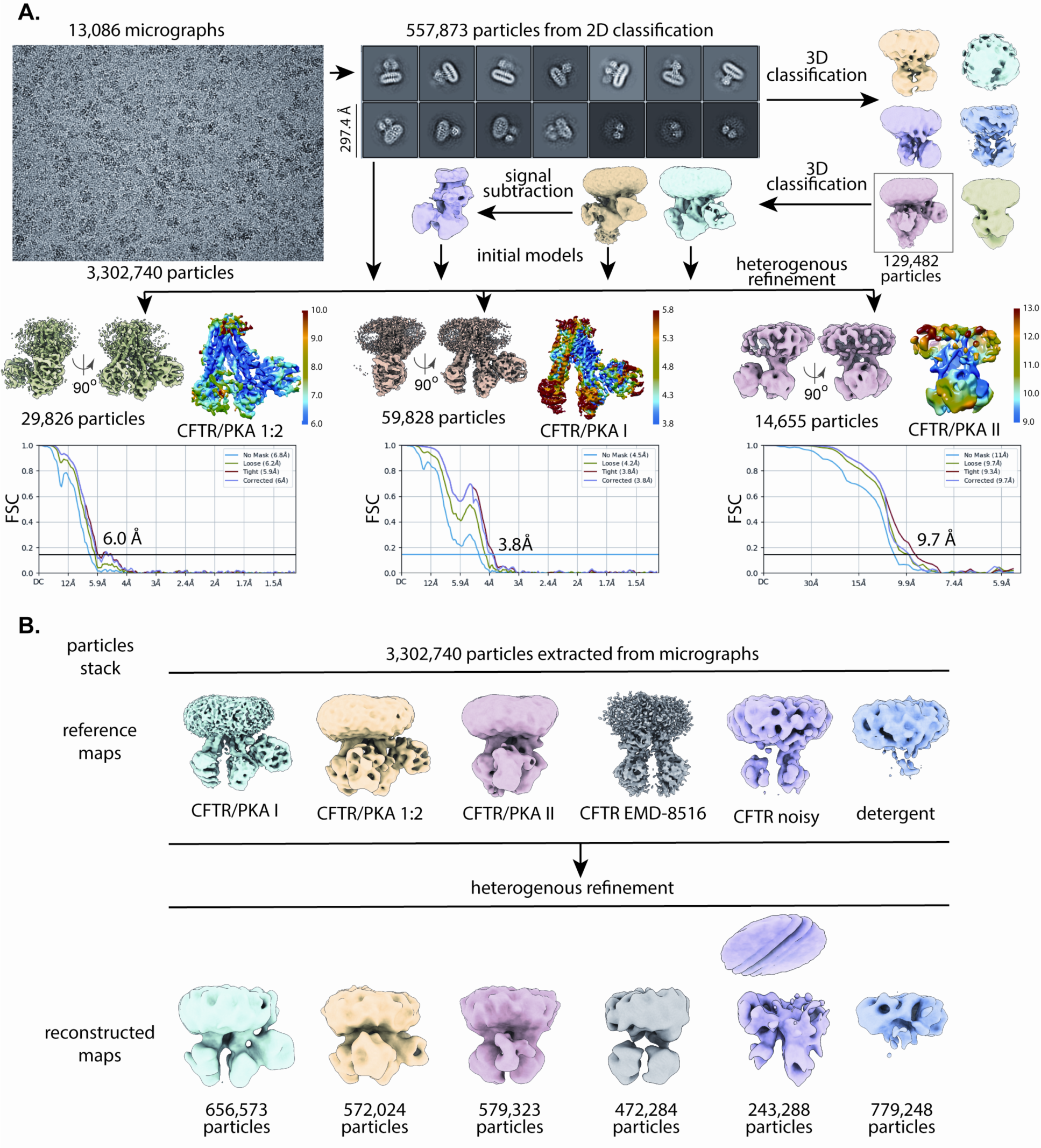
Cryo-EM reconstructions of PKA-C in complex with dephosphorylated CFTR. (A) Summary of image processing procedures and estimated resolution of each structure based on Fourier shell correlation (FSC) curves. (B) Heterogenous refinement of the entire dataset. Six structures were used as references for classification.

**Figure S2.**
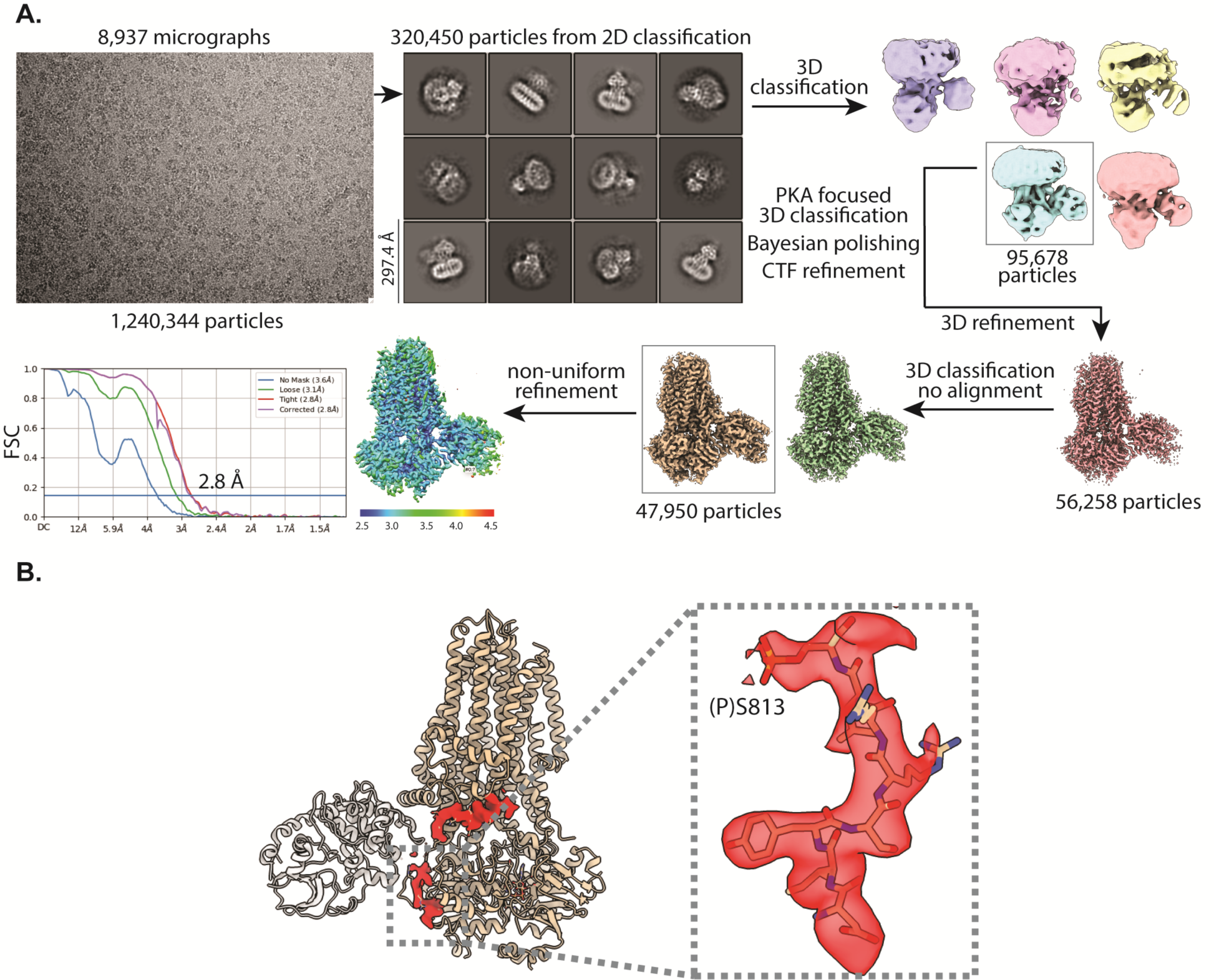
Cryo-EM reconstructions of PKA-C in complex with phosphorylated CFTR. (A) Image processing procedures and estimated resolution of PKA-C in complex with fully-phosphorylated, ATP-bound CFTR (E1371Q). (B) Density of residues 806-814 and 820-833 of the R domain.

**Figure S3.**
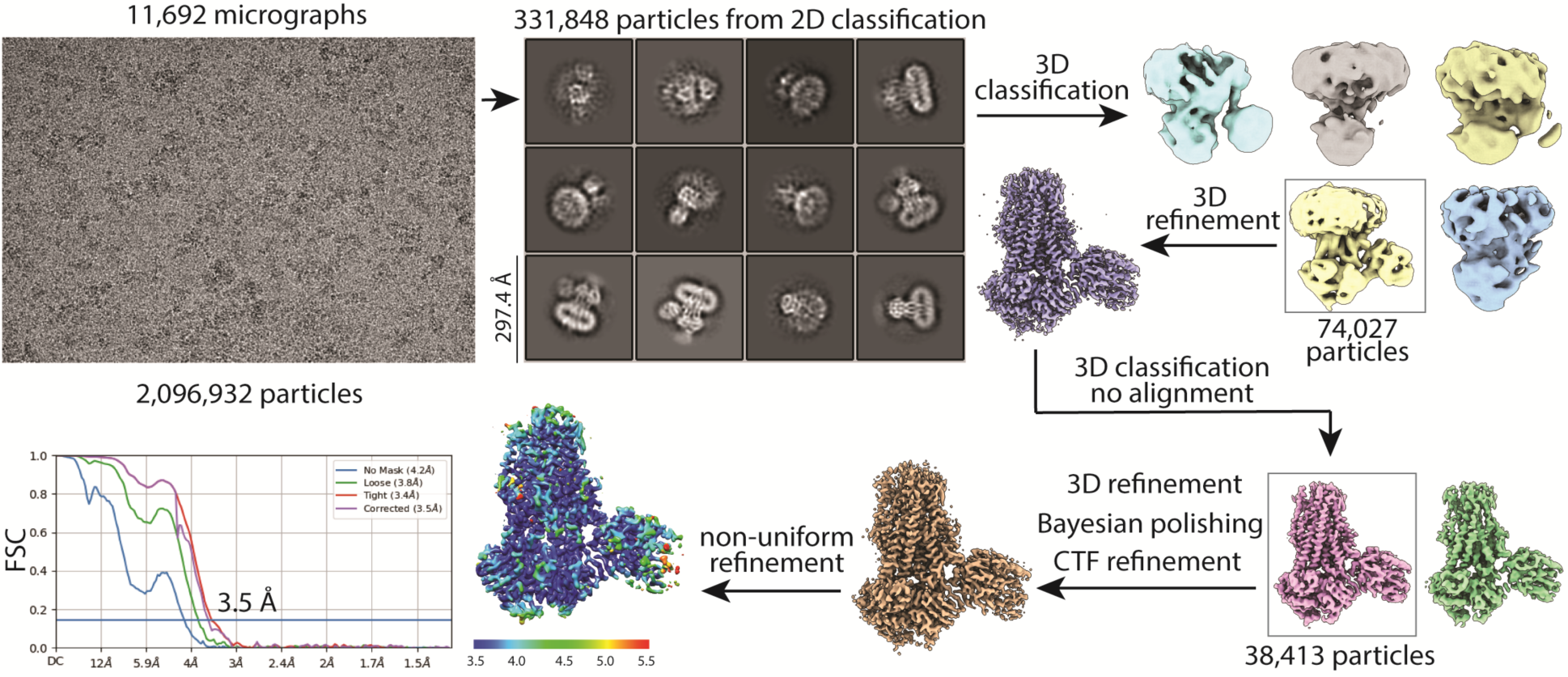
Cryo-EM reconstructions of PKA-C in complex with dephosphorylated CFTR, in the presence of P-ATP. Image processing procedures and estimated resolution of PKA-C in complex with dephosphorylated, ATP-bound CFTR (E1371Q) in the presence of P-ATP + AMPPNP.

**Figure S4.**
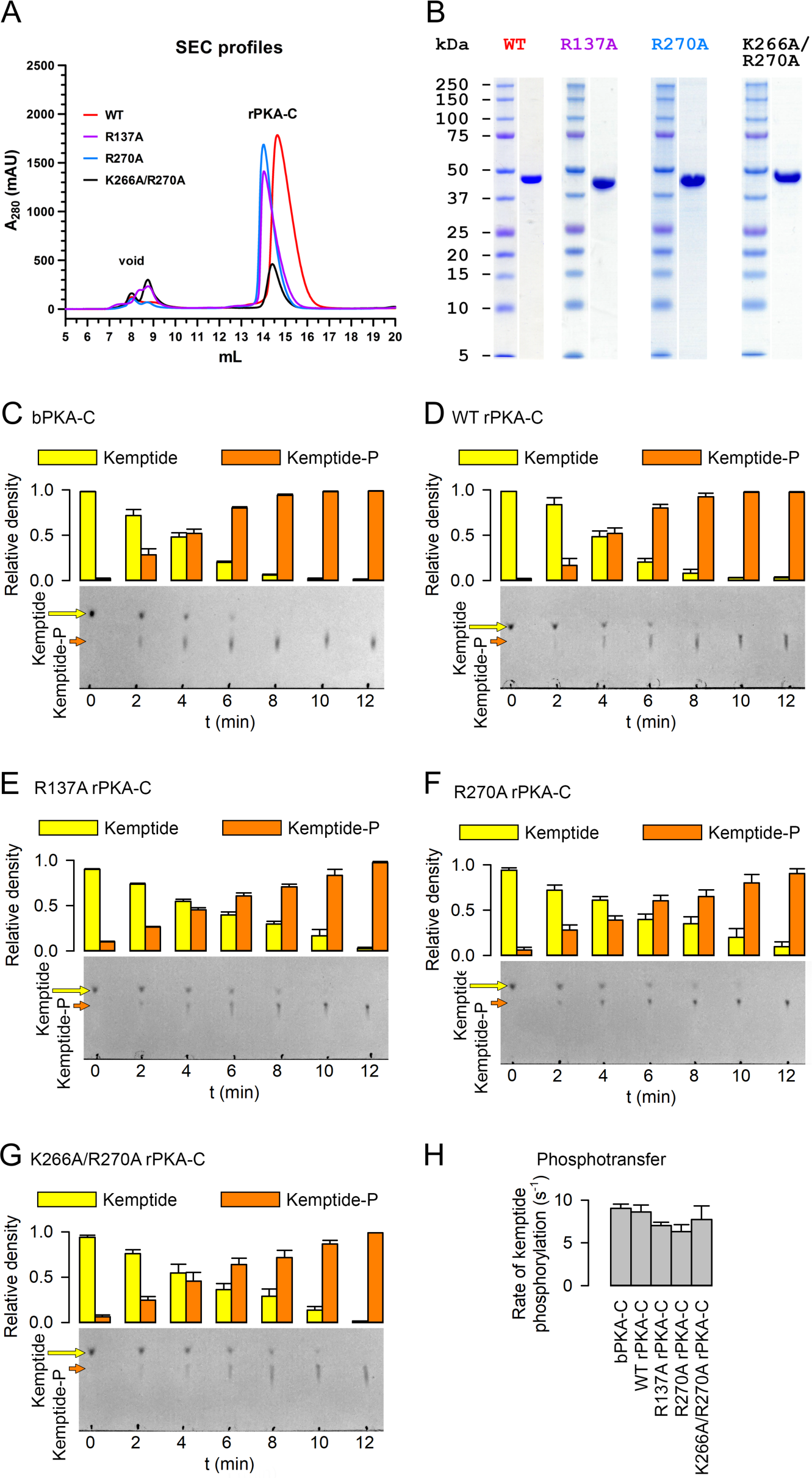
Purification of recombinant PKA-C (rPKA-C) constructs and rates of kemptide phosphorylation. (A) Size-exclusion chromatography (SEC; Superdex 200 10/300 GL, GE Healthcare Hungary) elution profiles, detected as absorbance at 280 nm, of WT (*red*), R137A (*violet*), R270A (B) (*blue*), and K266/R270A (*black*) rPKA-C. (B) Coomassie-stained SDS PAGE gels of the final pooled fractions from (A). Molecular weights for marker ladder bands (Precision, Bio-Rad) are labeled in kDa. (C)-(G); Time courses of kemptide phosphorylation resolved on TLC sheets (*Bottom*) and densitometric analysis (*Top*) for (C) bPKA-C, (D) WT rPKA-C, (E) R137A rPKA-C, (F) R270A rPKA-C, and (G) K266A/R270A rPKA-C. In each case 5 nM enzyme was incubated with 20 μM TAMRA-kemptide + 200 μM MgATP for the indicated amounts of time at room temperature, and 2 μl aliquots were spotted on the TLC sheets. (H) Calculated *k*_cat_ (s^-1^) for the five PKA-C proteins in (C)-(G). Data in (H) represent mean±SEM, n=3.

**Figure S5.**
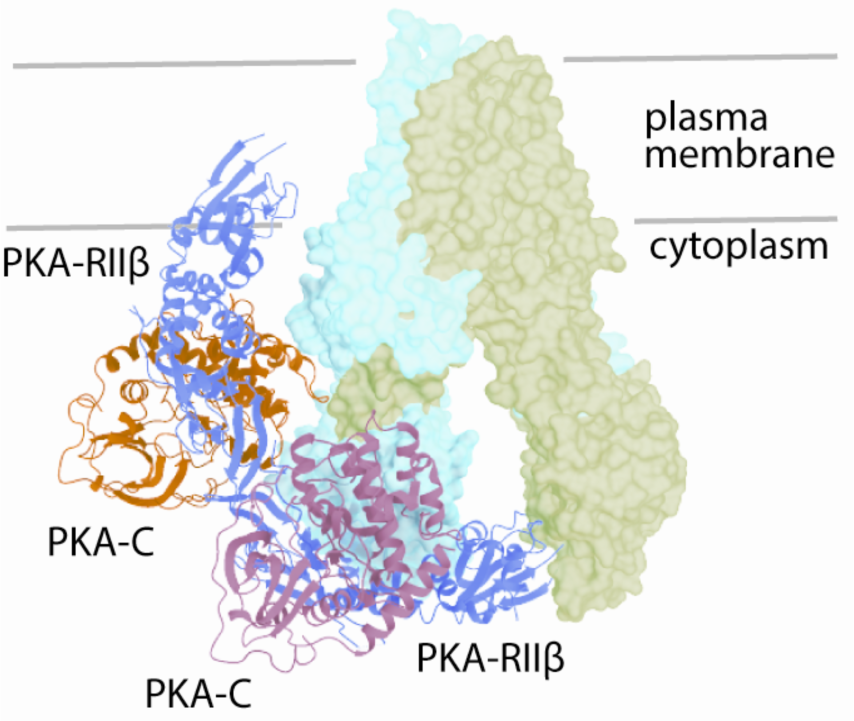
Alignment of PKA holoenzyme structure with CFTR-PKA _I_ complex. Structure of dephosphorylated CFTR-PKA_I_ complex aligned with the structure of the PKA (Cα)_2_(RβII)_2_ holoenzyme (PDBID: 3TNP). CFTR is shown as surface, PKA_I_ as orange ribbon. For the PKA holoenzyme (ribbon) the R subunits are shown in blue, the C subunits in purple. One C subunit of the holoenzyme was aligned with PKA_I_.

